# Enhanced Structural Brain Connectivity Analyses Using High Diffusion-weighting Strengths

**DOI:** 10.1101/2024.10.02.616308

**Authors:** Leyao Yu, Adeen Flinker, Jelle Veraart

**Affiliations:** New York University Tandon School of Engineering; New York University Grossman School of Medicine; Center for Advanced Imaging Innovation and Research (CAI2R), Department of Radiology, New York University Grossman School of Medicine

## Abstract

Tractography is a unique modality for the in vivo measurement of structural connectivity, crucial for understanding brain networks and neurological conditions. With increasing *b*-value, the diffusion-weighting signal becomes primarily sensitive to the intra-axonal signal. However, it remains unclear how tractography is affected by this observation. Here, using open-source datasets, we showed that at high *b*-values, DWI reduces the uncertainty in estimating fiber orientations. Specifically, we found the ratio of biologically-meaningful longer-range connections increases, accompanied with downstream impact of redistribution of connectome and network metrics. However, when going beyond *b*=6000 s/mm^2^, the loss of SNR imposed a penalty. Lastly, we showed that the data reaches satisfactory reproducibility with *b*-value above 1200 s/mm^2^. Overall, the results suggest that using *b*-values above 2500 s/mm^2^ is essential for more accurate connectome reconstruction by reducing uncertainty in fiber orientation estimation, supporting the use of higher *b*-value protocols in standard diffusion MRI scans and pipelines.

## Introduction

The brain is organized as complex networks of cortical and subcortical neurons connected via long-range myelinated axonal pathways. The architecture of neuronal networks is central to human cognition and the physiology of various diseases and disorders, as well as development and aging (***Glasser et al., 2016***; ***Douaud et al., 2014***; ***Tononi et al., 1994***; ***Park and Friston, 2013***). For example, language, a uniquely human ability, has been found to involve a network spanning inferior frontal, inferior parietal, and superior temporal cortices, linked by the arcuate fasciculus, a major tract critical to reading (***Yeatman et al., 2011***), semantics and comprehension (***Glasser and Rilling, 2008***), and underlies deficits in aphasia (***Bernal and Ardila, 2009***; ***Ivanova et al., 2016***). The emergence of neuroimaging methods such as functional (fMRI) and diffusion-weighted magnetic resonance imaging (dMRI) advanced the study of functional and structural brain connectivity (***Behrens and Sporns, 2012***) and its relation to cognition, in health and disease (***Fornito et al., 2015***).

Today, diffusion MRI is a unique technology for studying the intricate network of connections between the human brain in vivo (***Sotiropoulos and Zalesky, 2019***; ***Shamir and Assaf, 2024***). The dMRI signal is sensitive to interactions between diffusing protons and the microstructural environment of brain tissue at a length scale well below spatial resolution (***Bihan, 1986***; ***Beaulieu, 2002***). Diffusion MRI can be used for the development of biomarkers of tissue integrity (***Stanisz et al., 1997***; ***Novikov et al., 2018***) and tractography (***Mori and Van Zijl, 2002***). In particular, tractography is the digital reconstruction of fiber pathways through the long-range integration of localized fiber directions (***Jeurissen et al., 2019***). The technique is routinely used in preoperative planning for tumor resection (***Essayed et al., 2017***; ***Yeh et al., 2021***), epilepsy surgery (***Duncan et al., 2016***), neurosurgical procedures such as deep brain stimulation (***Calabrese, 2016***), the diagnosis of acute ischaemic stroke (***Moseley et al., 1990***), and safety monitoring of patients after drug therapy (***Wattjes et al., 2015***). Alongside its increasing clinical use, the wide adoption of dMRI in large-scale neuroimaging collections in neuroscientific and biomedical research poses great promise to the study of the organization of brain networks and their association with brain function at the population level (***Horien et al., 2021***; ***Schilling et al., 2023***; ***Elliott et al., 2018***; ***Smith et al., 2020***).

Despite wide application, the interpretation of structural connectivity from tractography remains challenging due to several limitations (***Schilling et al., 2019***; ***Maier-Hein et al., 2017***; ***Jbabdi and Johansen-Berg, 2011***). These include the lack of directionality, the low accuracy in determining in which cortical layers cortical-cortical connections terminate, the systematic observation of spurious connections, and missing connections (***Jeurissen et al., 2019***; ***Girard et al., 2014***). While some of these limitations are intrinsic to the technique, others might be addressed by recent advances in hardware and biophysical modeling(***Jones, 2010***; ***Afzali et al., 2021***). Indeed, improvement in spatial resolution, signal-to-noise ratio, and increased *b*-value are presumed to reduce the uncertainty in the estimation of local fiber directions (***Veraart et al., 2016***; ***Tournier et al., 2013***; ***Setsompop et al., 2013***). These advancements may improve the sensitivity of tractography to complex white matter configurations, and improve the identification of cortico-cortical connections, allowing for a better understanding of their interaction with the functional specialization of related networks.

Tournier et al. (***Tournier et al., 2007***) previously showed that the angular resolution of the fiber orientation distribution function increases with the *b*-value, thus promoting the use of *b*-values of minimally 3000 s/mm^2^ for tractography. More recently, studies suggest that at even higher *b*-values (i.e. 6000 s/mm^2^ and higher), the dMRI signal originates predominantly in the axons (***Veraart et al., 2020***). Although the use of such high *b*-values results in significant signal loss, leading to a lower signal-to-noise ratio (SNR), it is favorable for the accuracy of biophysical modeling, axon diameter mapping in particular (***Veraart et al., 2021, 2020***). Its impact on tractography and structural connectivity analysis remains underexplored. We hypothesize that the enhanced sensitivity for intra-axonal signal, relative to extra-axonal signal, holds the potential for more precise and accurate fiber tractography with downstream impact on structural network and network metrics.

In this study, we used open-access data that span a wide range of *b*-values to characterize the effect of the *b*-value on brain tractography and network analyses. Our findings demonstrate that higher *b*-values significantly enhance the robustness of structural networks and tractography, particularly in reconstructing longer-range connections.

## Methods and Materials

### Subjects and Data

We performed a secondary analysis of the previously published and publicly available *MICRA* data set. All details on subjects and imaging protocols are described in ***Koller et al. (2021)***. A summary is given here. Six research volunteers were scanned five times, using identical protocols, using a 3T Connectom MR scanner (Siemens). During each scan session, T1-weighted MPRAGE and multishell dMRI images were recorded. Diffusion weighting was applied with *b* = 200, 500, 1200, 2400, 4000, and 6000 s/mm^2^. The number of gradient directions are 20, 20, 30, 61, 61, and 61 for these *b*-shells. Furthermore, 11 *b* = 0 images were recorded interspersed. The following scan parameters were kept constant: TR/TE= 3000/59ms, matrix=110 x 110, and 62 slices with a corresponding voxel dimension of 2×2×2 mm^2^.

### Preprocessing

All data were preprocessed using a unified pipeline that includes: denoising, Gibbs ringing correction, and correction for geometric distortions due to subject motion, eddy current, and magnetic susceptibility effects similar to ***Ades-Aron et al. (2018***). All corrections were performed using MR-trix3.0 tools on the BrainLife platform (***Hayashi et al., 2024***).

### Cortical parcellation

For each subject, cortical parcellation was derived from high-resolution T1-weighted MPRAGE images using the FreeSurfer pipeline (***Fischl, 2012***). The Desikan-Killiany atlas was adopted to define 68 cortical nodes for the network analysis (***Desikan et al., 2006***). The T1-weighted images were rigidly registered to the diffusion imaging data after the parcellation and rigid transformation is applied to such parcellation maps. The subcortical structures and the cerebellum were omitted from the analysis.

### Tractography

Whole-brain, probabilistic tractography was performed using anatomical constraints. The Freesurfer output image was used to define white matter, gray matter, and cerebrospinal fluid. The MRtrix3.0 package is used for fiber orientation estimation and tractography using iFOD2 with the default settings (***Tournier et al., 2019***). Unless specified otherwise, all tractography was seeded and terminated at the white and gray matter interface.

### Quantifying uncertainty in orientation coherence

We used bootstrapping to calculate the uncertainty of fiber orientation estimates as a function of the *b*-value. For each bootstrap realization (*N*=500), we selected a random sample of repeated measurements. All bootstrap realizations were unique because of the random drawing of individual data points from one out of five repetitions, but they ultimately had identical diffusion encoding schemes. For each bootstrap realization and each *b*-value, we estimated the single-fiber signal kernel, computed the fODF using constrained spherical deconvolution, and derived the peak of the main fiber direction using MRtrix3.0 tools (***Tournier et al., 2019***). Per WM voxel, we measured the variability of the principal diffusion peaks using a coherence metric that was derived from the eigenvalues of the dyadic tensor (***Jones, 2003***). In addition, we computed the 95^th^ percentile in the angular error to the average peak. Similar to ***Veraart et al. (2016***), we avoided a sorting bias by selecting the primary peak orientation based on their alignment with the primary peak of the fODF that was obtained by averaging all bootstrap realizations.

### Network Analysis

The 68 × 68 connectivity matrices, derived from Desikan-Killiany atlas (***Desikan et al., 2006***), were constructed by counting the number of streamlines that connect to corresponding cortical areas and measuring their average length. Commonly used graph-theoretical metrics (average network degree and strength, betweenness, centrality, clustering coefficient, and normalized global efficiency) were calculated using iGraph package (***Csardi and Nepusz, 2006***; ***Csárdi et al., 2024***).

### Quantification of streamline length

We measured both the distance and displacement of streamlines. Both metrics are a measurement of length. However, the distance measures the length between the start- and end-point of a streamline along the curved sections of the streamline, whereas the displacement is the Euclidean distance between the start- and endpoint (Figure 2A). These length metrics are derived for each streamline but can be synthesized per node or nodal pair.

**Figure 1.**
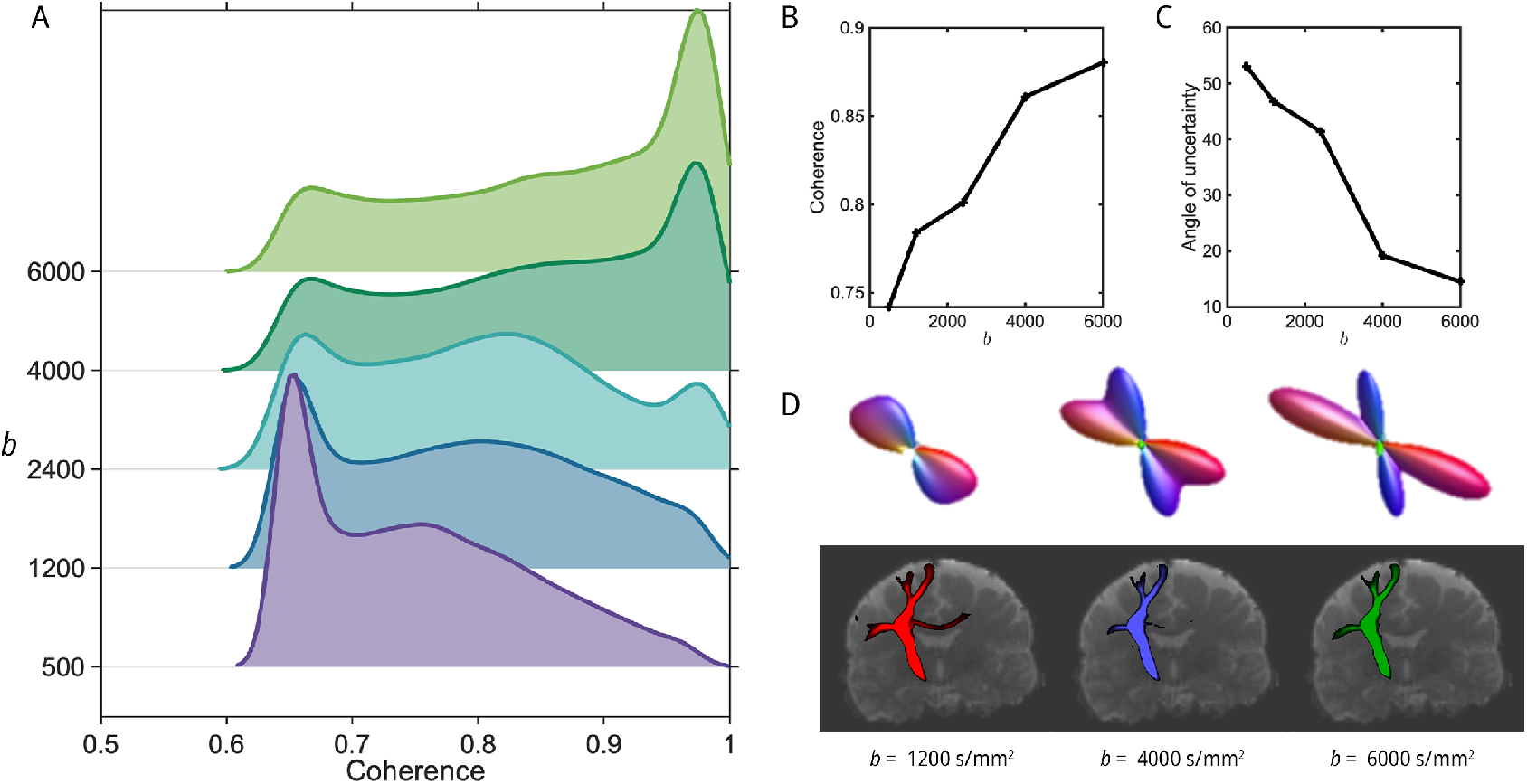
(A) The probability density function of orientation coherence in the estimation of the largest peak of the fODFs in all WM voxels for all *b*-values. The coherence was computed voxel-wise using 1000 bootstrap estimates. (B) The median of the orientation coherence distribution is shown as a function of *b*-value. (C) The 95% Confidence Intervals in the estimation of the fiber orientation (largest peak) is shown as a function of *b*-value (D) Top: exemplary fODFs are shown for various *b*-values. All fODFs were extracted from the same voxel, presenting a crossing fiber. Bottom: corticospinal tract for same set of *b*-values.

**Figure 2.**
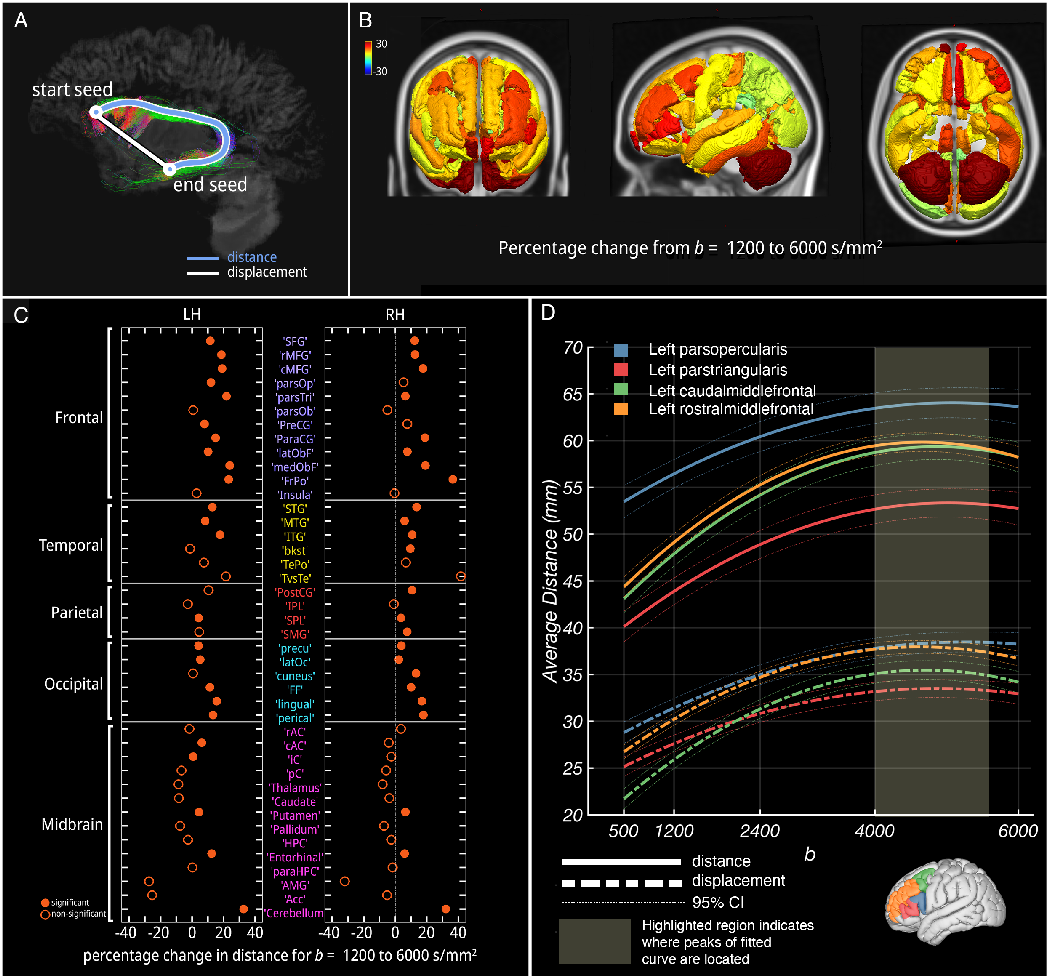
(A) Analysis paradigm depicting quantifying distance (travel distance) and displacement (Euclidean distance) between nodes. (B) Degree of change for average streamline length (travel distance) of each node from *b* =1200 to 6000. (C) Change in travel distance for each node from *b* =1200 to 6000 s/mm^2^. Solid circles stand for significant model fit. (D) Example of distance, displacement, and model fit of four critical language regions in left dorsolateral prefrontal cortex (Inferior Frontal Gyrus: pars triangularis, pars opercularis, Middle Frontal Gyrus: caudal and rostral middle frontal). Notable increment was observed with increasing *b* and peaked around *b* = 4000 s/mm^2^. The shaded area indicates vertices of model fit.

### Quantification of streamline density

For a pair of cortical nodes, *i* and *k*, we computed a streamline density as log 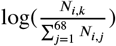 with *N*_*i*,*j*_ the number of streamlines connecting the *ith* and *jth* cortical. Per subject, the streamline densities were averaged across repetitions to improve precision.

### Statistical analysis

#### Coefficient of variation

For each cortical node, network metric, and *b*-value, coefficient of variation, CV, was calculated as 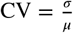, where *σ* is the standard deviation across the repeats, and *μ* is the mean of the repeats.

#### Polynomial mixed-effects model

To probe the effect of *b* on a given measure, we fit a polynomial mixed-effects model (matlab r2024a, fitlme function). We follow the model as such *y*_*ij*_ = *β*_0_ +*β*_1_ *⋅* (*b*-value_*ij*_)^2^ +*β*_2_ *⋅ b*-value_*ij*_ +*u*_*j*_ +*∈*_*ij*_, where *y*_*ij*_ is the response variable for the *i*_*th*_ observation in the *j*_*th*_ subject, *u*_*j*_ is the random effect for the *j*_*th*_ subject, and *∈*_*ij*_ is the residual error.

## Results

### Uncertainty in fODF estimation

We first report the impact of *b*-value on fiber orientation distribution functions (fODF). We found that a higher *b*-value results in higher orientation coherence across all WM voxels (Fig. 1A, 1B), and lower angle of uncertainty (Fig. 1C). The example of fODFs extracted from the same voxel with crossing fibers (Fig. 1D top) shows a clearer separation of different fiber bundles with a higher *b*-value. We highlighted the corticospinal tract by anatomical seeding and demonstrated the down-stream impact of fODF estimation on fiber tracking(Fig. 1D bottom). Despite the use of identical ROIs and tractography settings, the resulting tract varies widely between the evaluated *b*-values, with more spurious and commissural fibers at lower *b*-values.

### Impact of *b*-value on the length of streamlines

Next, we quantified the impact of *b*-value on the length of streamlines from a network perspective. All streamlines are seeded and terminated at the WM and GM interface and are labeled by the nodes to which they connect. We measured both the distance and displacement originating from each node (see Fig. 2A). We observed that both distance and displacement increase as a function of *b*-values in a region-specific basis (Fig. 2B, 2C). Most of the significant increments in distance are from neocortical regions such as bilateral frontal, temporal, and occipital cortices. Focusing on critical regions for the language network, we found that the average length of streamlines originating from Inferior Frontal Gyrus (IFG) and Middle Frontal Gyrus (MFG) significantly increased by 24.08% and 34.70% in the same *b*-range (Fig. 2D), although there is convergence around *b* = 4000 s/mm^2^. The observation that the distance and displacement change in a concordant way implies certain nodes form more long-range connections in higher *b*-values. Additionally, we observed a decrease in streamline distance between fixed node pairs as *b*-values increased (See Supplementary Figure 1).

### Impact of *b*-value on structural connectivity

Next, we evaluated whether the changes in streamlines have a downstream impact on the structural connections within brain networks. We focused on the language network, with pars triangularis, a node with known connections to the temporal cortex via the arcuate fasciculus, as well as its close association with neighboring nodes like pars opercularis. Overall, we found that the structural connectivity of pars triangularis changes from clinical standard to higher *b*-value, but remains stable when *b*-values are beyond clinical regime (Fig. 3A). Specifically, the structural changes are distinct to the location and innate structural connectivity between nodes. For example, the spurious short-range connections between pars triangularis and a neighboring node (e.g. pars opercularis) significantly decreased when *b* is equal and greater than 2400 s/mm^2^ (Fig. 3B). In comparison, for the meaningful long-range nodal connections from pars triangularis to left superior temporal gyrus (Fig. 3C), streamline density was increased by 160.6% from *b* = 1200 to 6000 s/mm^2^. Moreover, spurious long-range connections (e.g. pars triangularis to right anterior cingulate) significantly decreased when *b* is greater than 1200 s/mm^2^ (Fig. 3D). Further, we examined the meaningful long-range connection increment by examining the interhemispheric connection density. The left precentral gyrus connection to its right counterpart increased by 235.9% (Fig. 3E, 3F). Taken together, *b*-values beyond clinical standard provide more biologically plausible structural connectivity and less spurious connections, both short- and long-range.

**Figure 3.**
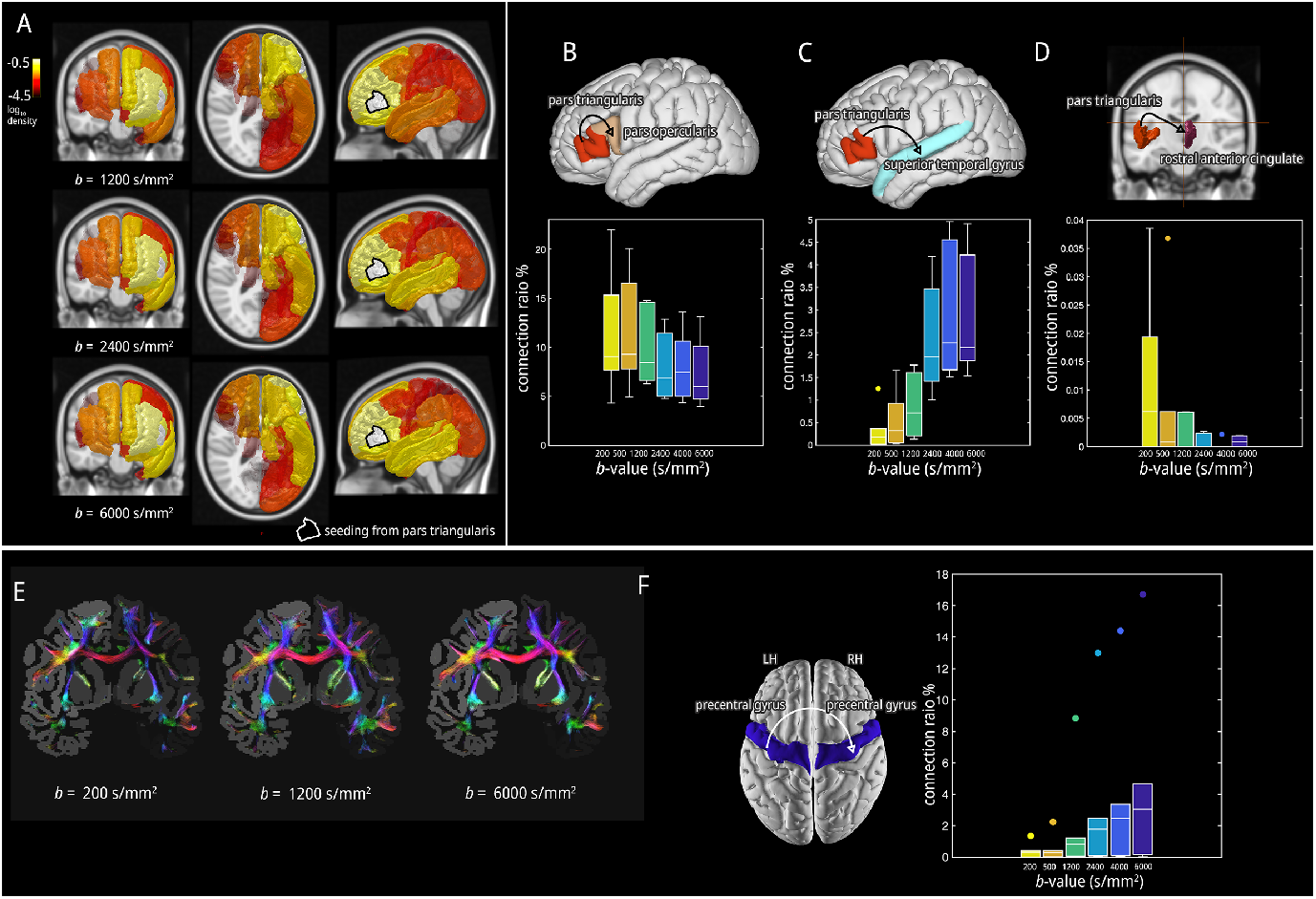
(A) degree of change in streamline density from pars opercularis to other nodes. Color is shown in *log*10 scale. Higher *b* shows higher density to cross-cortical regions. (B) Visualization of corpus callosum with increasing *b*. (C) Quantification of increment in streamline density from pars opercularis to superior temporal gyrus (STG). (D) Quantification of decrement in streamline density for nearby regions (pars opercularis to pars triangularis). (E) Increment in streamline density for cross-hemisphere connections from left precentral gyrus to right precentral gyrus.

### Impact of *b*-value on network metrics and reproducibility

Lastly, observing the changes in streamline distributions, we anticipated that the *b*-value would have downstream effects on the whole-brain connectome. To capture these effects, we analyzed changes in network metrics. Global changes in network strength is limited to 3.71%, while certain regions are more vulnerable to *b*-value changes. In bilateral superior frontal gyrus and other subcortical areas, the network strength change dramatically (SFG 23.20%, thalamus 21.36%, and cerebellum 52.83%, see Fig. 4A, 4B).

**Figure 4.**
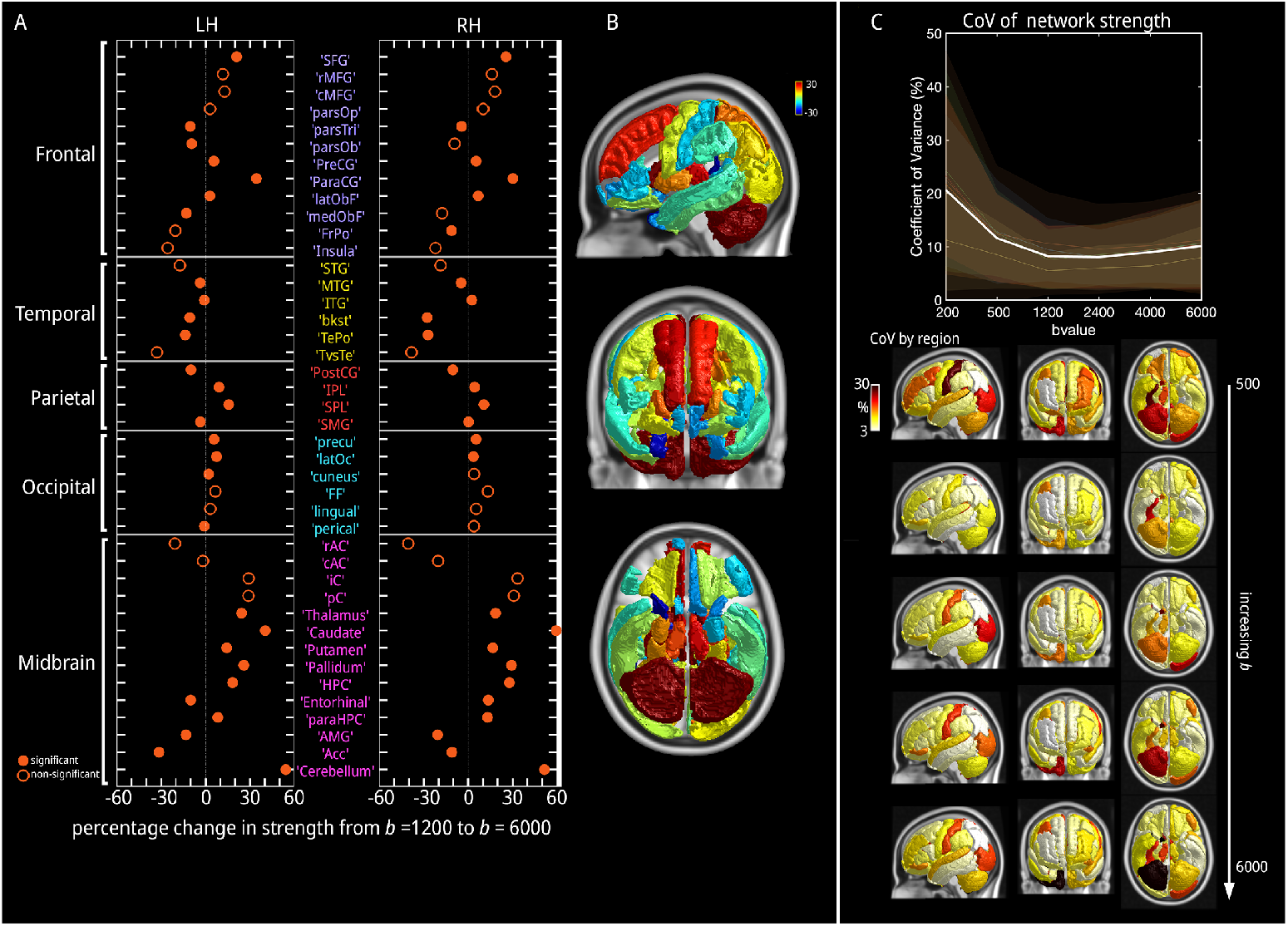
(A) Left: change in nodal strength from *b* = 1200 to 6000 s/mm^2^ for each node. Solid circles indicate a significant model fit. (B) visualization of the degree of change on the cortex. (C) Top: coefficient of variation (COV) across *b* for network strength. White lines stand for the mean across all subjects. Bottom: projection of COV for each node across increasing *b*.

To quantify reproducibility, we calculated the coefficient of variation (CoV) for global and local network metrics. For example, we found the CoV of network strength remains around 10% when *b* equals and greater than 1200 s/mm^2^. Using ANOVAN analysis, we do not observe loss in reproducibility of the findings for *b*-values above 1200 s/mm^2^, despite the lower SNR at high *b*-values (Fig. 4C).

## Discussion

In this study, we evaluated the impact of *b*-values on tractography and structural connectivity analyses. Existing studies have not adequately captured the impact of high *b*-values in the living human brain. To address this limitation, we established a framework to analyze dMRI data with repeated measures, on which tractography and post-analysis were performed. Specifically, we evaluated the changes in fODF, streamline length, tractography, local network, graph measures, and reproducibility measures with *b*-values. Our primary finding is that the proportion of large-range connections between distinct cortical nodes increases, while spurious short-range connections decrease with increment in *b*-value.

Historically, diffusion MRI data have been acquired with *b*-values in the range of *b* = 800 - 1200 s/mm^2^. Such *b*-values are optimal for diffusion tensor imaging in the living human brain. Today, many neuroimaging studies and clinical scan protocols still adopt such practices. However, there is a notable trend to acquire more and higher *b*-values to promote biophysical modeling and to improve the angular resolution for tractography (***Tournier et al., 2013***). For example, various research consortia use different *b*-values in their diffusion MRI protocols, ranging from 500 to 3000 s/mm^2^ (***Casey et al., 2018***; ***Jack et al., 2008***; ***Miller et al., 2016***; ***Taylor et al., 2017***; ***Van Essen et al., 2013***). However, recent developments in imaging hardware, i.e. ultra-gradient gradients, now allow for the acquisition of even higher *b*-values, (i.e. *b*=6000 s/mm^2^ and up), with little penalty on echo times and the signal-to-noise ratio (***Setsompop et al., 2013***; ***Fan et al., 2016***). At higher *b*-values, the diffusion-weighted signal becomes more sensitive to intra-axonal processes, though at the cost of significant signal loss. Yet, the impact, or confounding factors, of the use of such high *b*-values on diffusion MRI processing and analysis requires further study.

Alongside other experimental factors, e.g. spatial resolution and echo time, the *b*-value has a quantitative impact on diffusion MRI metrics, including DTI and DKI (***Qin et al., 2009***; ***Veraart et al., 2011***). In the absence of harmonized imaging protocols, the *b*-value dependency of diffusion metrics is a culprit in the reproducibility, comparability, and interpretability of studies (***Cetin-Karayumak et al., 2024***). Yet, at the same time, protocols can be optimized in terms such as experimental factors to promote the accuracy and precision of downstream analyses. For example, previous studies have shown that the use of higher *b*-values results in an increased angular resolution, “sharper” fODFs, increased ability to disentangle crossing fibers, and, more generally, promote more accurate fiber tractography (***Tournier et al., 2013***; ***McKinnon et al., 2017***; ***DeLano et al., 2000***; ***Hui et al., 2010***; ***Fan et al., 2016***; ***Xie et al., 2015***). In our study, we complement such studies by evaluating the impact of the *b*-value on reconstructed streamlines and structural network metrics.

Previously, the *b*-value dependency of the structural connectome was evaluated in postmortem rodent brains (***Crater et al., 2022***). The study found that a high *b*-value around 3000 s/mm^2^, with a single-shell acquisition of 384 gradients, was sufficient to resolve crossing fibers, resulting in a connectome that most closely resembled the ground truth. In contrast, higher *b*-values (8000 s/mm^2^) led to an increase in false positive streamlines.

Here, we reproduced these findings demonstrating reduced uncertainty in the estimation of fiber directions with increasing *b*-values, leading to a notable reduction of spurious fibers in tractography. Our findings that the decrease of angle of uncertainty and increase of coherence in (Fig. 1 aligned with the findings in (***Xie et al., 2015***; ***Tournier et al., 2013***). ***Xie et al. (2015***) observed that fODF peaks became sharper at higher *b*-values, with *b*-values of 2000 s/mm^2^ and above revealing most two-way or three-way crossing-fiber structures. We showed that the coherence continues to increase beyond the clinical regime (i.e. 2500 s/mm^2^). We also expanded our findings with ***Tournier et al. (2013***)’s findings on single fibers, where they identified *b*-value = 3000 s/mm^2^ and l = 8 as providing the highest achievable angular resolution. On the topic of tract reconstruction, our results in Fig. 1D showed a similar trend as in***Fan et al. (2016***), where fODF reconstruction at the intersection of the corticospinal tract and corpus callosum was visually sharper with each increasing *b*-value up to *b* = 10000 s/mm^2^.

The impact of *b*-value was extended to anatomically-constrained tractography. Our results are consistent with previous studies noting a global increase in streamline distance with higher *b*-values (***Khalilian et al., 2021***). Additionally, we identified a statistically significant increase in streamline distance within specific nodes, indicating that this effect is region-specific rather than global. This change in streamline length may arise from three potential factors: (1) more wandering streamlines in existing connections, (2) an increase in existing long-range connections, or (3) the formation of new connections to distant nodes. If the first scenario were true, we would expect to observe longer streamline distance between fixed node pairs. However, we found that, on average, streamline distance between fixed node pairs was shorter at higher *b*-values (Supplementary Fig. S1A).

Our data supports the second scenario (2), where the density of long-range connections increased within fixed node pairs (Supplementary Fig. S1B). For instance, in node pairs within the language network (cMFG, rMFG, pars triangularis, pars opercularis, STG, and MTG), we observed that the impact of *b*-value differed between short- and long-range connections. Higher *b*-value reduced the ratio of shorter connections while increasing the density of long-range, cross-cortical connections by one to two orders of magnitude.

The third scenario seems unlikely to contribute to the lengthening of streamlines. Upon inspecting the emergence and disappearance of connections between low and high *b*-values, we found that most changes involved the disappearance of connections, with subcortical structures being the most affected (Supplementary Fig. S1C). This is likely due to the increased SNR, which decreases the number of streamlines terminating in nearby subcortical GMWM interface. Additionally, higher *b*-value may introduce spurious long-range, cross-hemisphere connections. As shown in Supplementary Fig. S1A, some longer connections (above 150 mm) form between distant, cross-hemispheric node pairs, which may be attributed to streamlines that cross the corpus callosum and traverse other major association tracts. In summary, our findings support the second scenario, where higher *b*-values facilitate the formation of more precise long-range connections, leading to altered structural connectomes.

Following the redistribution of connectomes, we further observed that *b*-value also impacts network metrics. Consistent with previous literature, we find that the impact of *b*-value is region-specific (***Caiazzo et al., 2018***). Our analysis shows that the largest increase in network strength occurs primarily in subcortical regions, such as the caudate, thalamus, and cerebellum. Since network strength is measured by streamline counts, higher strength in these small subcortical regions may indicate that more streamlines are connecting to other anatomical regions rather than terminating early. In addition to the above findings, we demonstrate that the use of higher *b*-values is not associated with a drop in reproducibility. Our findings encourage the adoption of higher *b*-values in clinical settings and multi-modal studies without a loss in reproducibility.

In this study, we focused on the language network to demonstrate how higher *b*-values impact connection density, particularly in long-range connections like those between pars triangularis and superior temporal gyrus, connected via arcuate fasciculus (AF). The AF, with its complex fiber structure, is often susceptible to variations in imaging protocols, making it a challenging tract for precise reconstruction(***Catani and Mesulam, 2008***; ***Duffau, 2008***; ***Frey et al., 2008***). Our findings indicate that higher *b*-values reduce the uncertainty in fODFs in voxels where turning and crossing happen. This reduction in uncertainty improves the precision of tractography, allowing streamlines to terminate more accurately at their anatomical destinations. Our finding is in line with previous studies, such as ***Maffei et al. (2019***), which showed similar benefits of higher *b*-values in reconstructing acoustic radiation, with false positives significantly decreasing as *b*-values increased. The increased precision and reduced variability with higher *b*-values support future studies that combine diffusion imaging with other modalities, such as functional MRI or EEG, to explore the structural and functional connectivity of the language network more comprehensively.

In the current paper, we used anatomically constrained tractography to reduce the sampling bias (***Smith et al., 2012***). The method improves the biological accuracy of streamline connections by using anatomical information to control the evolution and termination of tractography. However, we did not evaluate tractography filtering techniques such as SIFT (***Smith et al., 2013***), SIFT2 (***Smith et al., 2015***), COMMIT (***Daducci et al., 2015***), or LiFE (***Pestilli et al., 2014***). Such techniques are known to affect structural network metrics such as global efficiency or clustering coefficient, while the number of false positive streamlines may be reduced at the expense of filtering true connections as well. Yet, the reproducible of reconstructing the connectomes has shown to be reduced for some of the filtering techniques, when compared to the unfilterd streamlines ***Koch et al. (2022***); ***Frigo et al. (2020***); ***Sarwar et al. (2023***).

The study presents a secondary analysis of the MICRA data sets. Within the scope of our work, the data prevents us from evaluating the *b*-value trend beyond *b* = 6000 s/mm^2^. Previously, it has been shown that *b* = 6000 s/mm^2^ is a tipping point in the respective contributions of the intra- and extra-cellular compartments to the dMRI signal. Indeed, at b-6000, the extra-cellular signal, more anisotropic in its nature, has been fully suppressed and the residual signal is presumed to be specific to axons, and possibly glial processes. We confirmed this trend by analyzing another publicly available HCP data set with *b*-values up to 10,000 s/mm^2^ (***Fan et al., 2016***) (Supplementary Fig. S2). The accuracy of tractography plateaued around *b* = 5000 s/mm^2^, and dropped at 10000, due to an increasing loss in the signal-to-noise ratio.

We limit our evaluation to anatomically constrained tractography following constrained spherical convolution of the dMRI signal with a *b*-value specific, yet data-driven response function. We used the implementation and default settings of the widely adopted MRtrix software library. The observed *b*-value dependency is likely to impact alternative tractography techniques in similar manners, yet a comparative analysis is beyond the scope of this study. Amongst the alternative strategies, multi-shell multi-tissue CSD has been presented to exploit the unique *b*-value dependencies of the different macroscopic tissue types (WM/GM/CSF) to improve the reliability of fiber tractography. However, the use of high *b*-values is a filter for WM, axons in particular, thereby mitigating the need for modeling the *b*-value dependency of the tissue-specific signal kernel.

## Acknowledgments

This work was supported by National Institute of Health grants P41 EB017183, R01NS088040, R01NS109367 (A.F.), and R01NS115929 (A.F.). The work was partially performed at the Center of Advanced Imaging Innovation and Research (CAI2R, www.cai2r.net), and NIBIB Biomedical Technology Resource Center.

## Data and Code availability

The data used in the analysis and results sections were acquired at the UK National Facility for In Vivo MR Imaging of Human Tissue Microstructure funded by the EPSRC (grant EP/M029778/1), and The Wolfson Foundation.

The data in Supplementary Fig. S2 were from the Human Connectome Project, WU-Minn Consortium (Principal Investigators: David Van Essen and Kamil Ugurbil; 1U54MH091657) funded by the 16 NIH Institutes and Centers that support the NIH Blueprint for Neuroscience Research; and by the McDonnell Center for Systems Neuroscience at Washington University.

The code is available upon publication at https://github.com/flinkerlab/.

## Declarations

J.V. is a co-inventor of patent US20180120404A1 (licensed) describing the denoising technology that is used in this study.

## Supplementary Information

**Figure S1.**
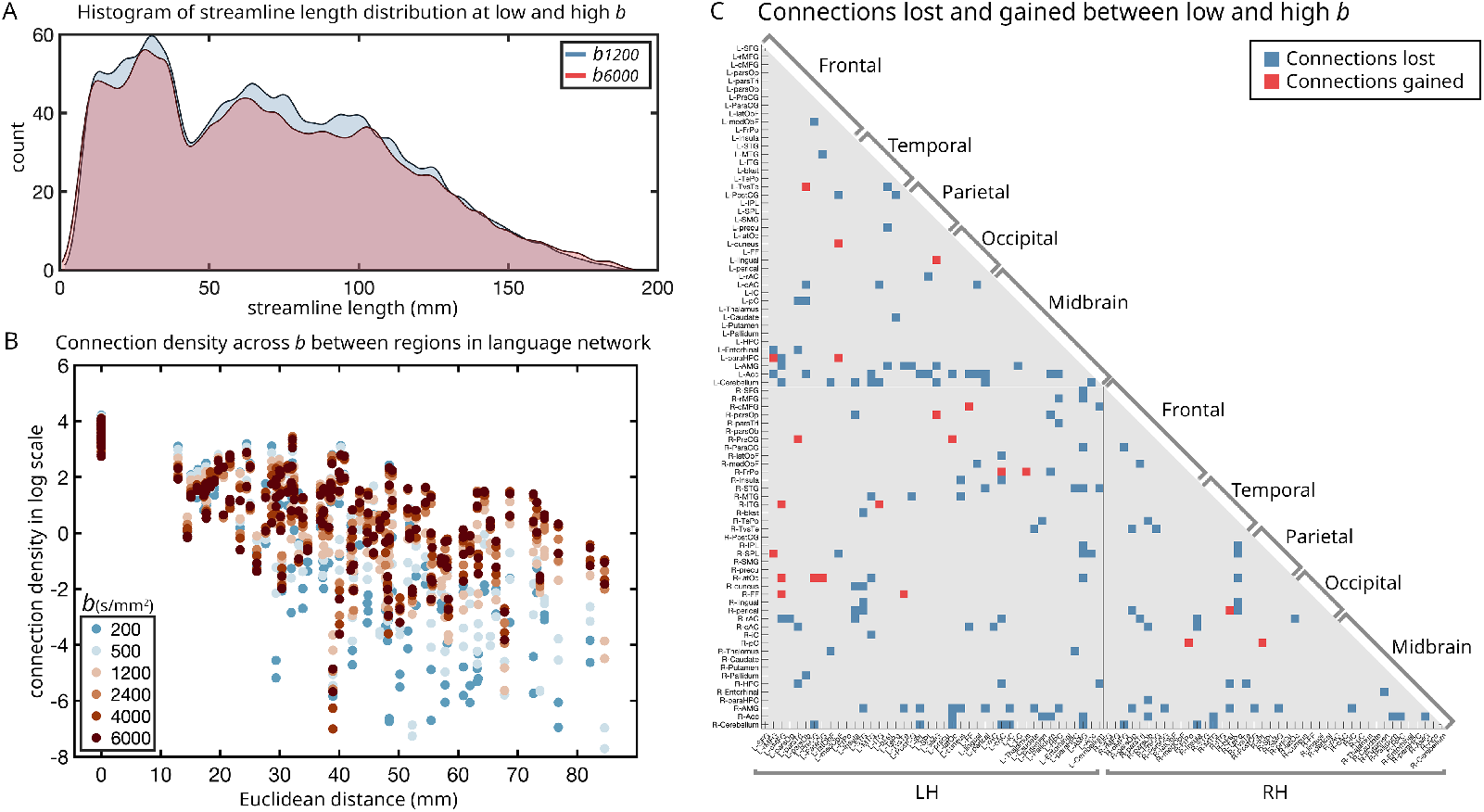
(A) Distribution of streamline length for fixed node pairs at *b* = 1200 and *b* = 6000 s/mm^2^. (B) Connection density across varying *b*-values between specific node pairs within the language network. Six anatomical regions (cMFG, rMFG, pars triangularis, pars opercularis, STG, and MTG) were selected as representative areas within the language network. The Euclidean distance between each node pair was derived from subject-specific anatomical centroids. Connection density (i.e., the percentage of streamline counts in the specified node pair divided by the total streamlines originating from that pair) is plotted on a logarithmic scale. (C) Emergence and disappearance of connections in the connectome. Changes are considered only if they occur in more than half of the subjects.

**Figure S2.**
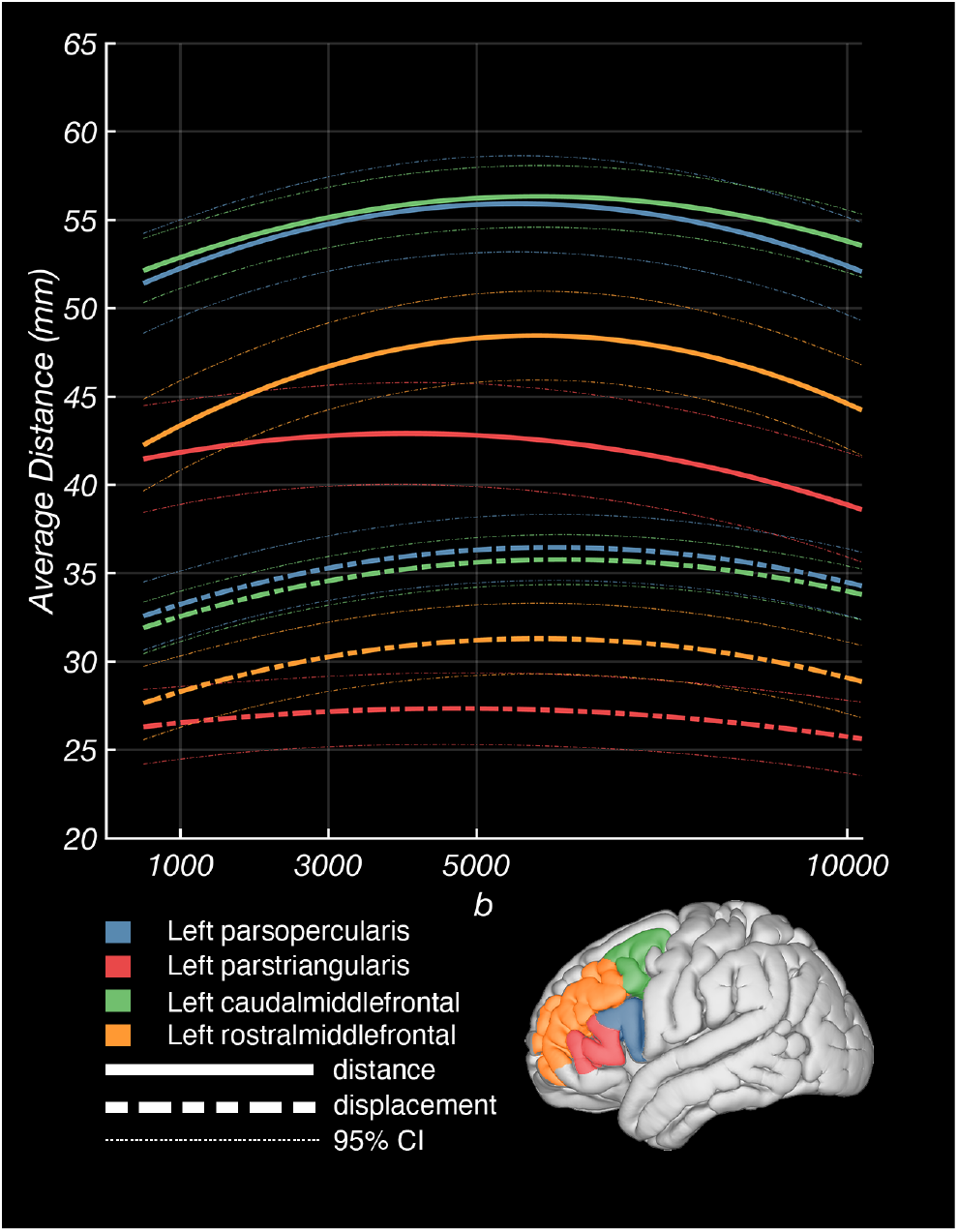
Example of distance, displacement, and model fit of four critical language regions in left dorsolateral prefrontal cortex for HCP ultra high **b**-value dataset.

