## SupplementaryMaterials for "Enhanced Structural Brain Connectivity Analyses Using High Diffusion-weighting Strengths"

### Supplementary Information

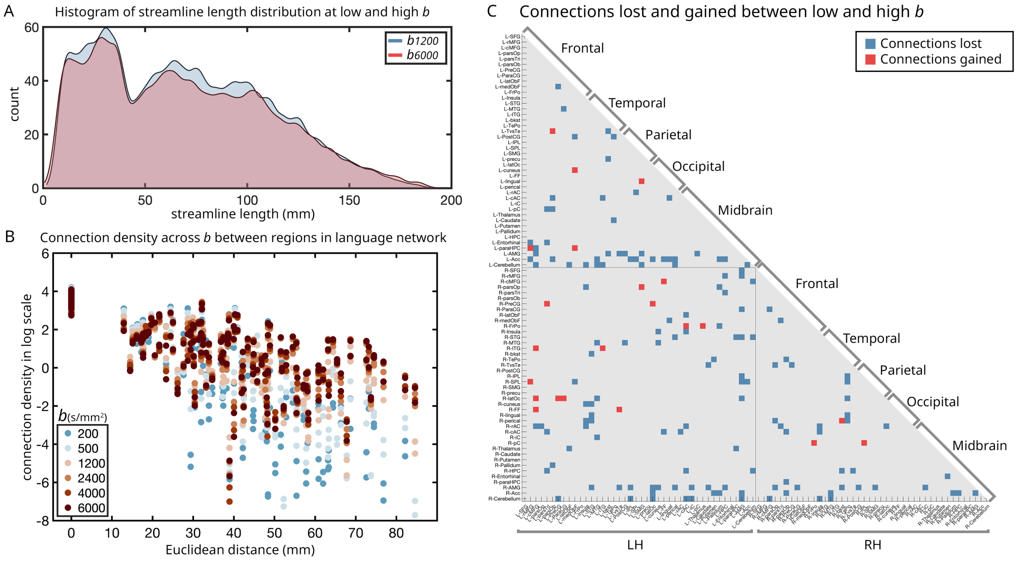


**Figure S1.** (A) Distribution of streamline length for fixed node pairs at *b* = 1200 and *b* = 6000 s/mm$^2$. (B) Connection density across varying *b*-values between specific node pairs within the language network. Six anatomical regions (cMFG, rMFG, pars triangularis, pars opercularis, STG, and MTG) were selected as representative areas within the language network. The Euclidean distance between each node pair was derived from subject-specific anatomical centroids. Connection density (i.e., the percentage of streamline counts in the specified node pair divided by the total streamlines originating from that pair) is plotted on a logarithmic scale. (C) Emergence and disappearance of connections in the connectome. Changes are considered only if they occur in more than half of the subjects.

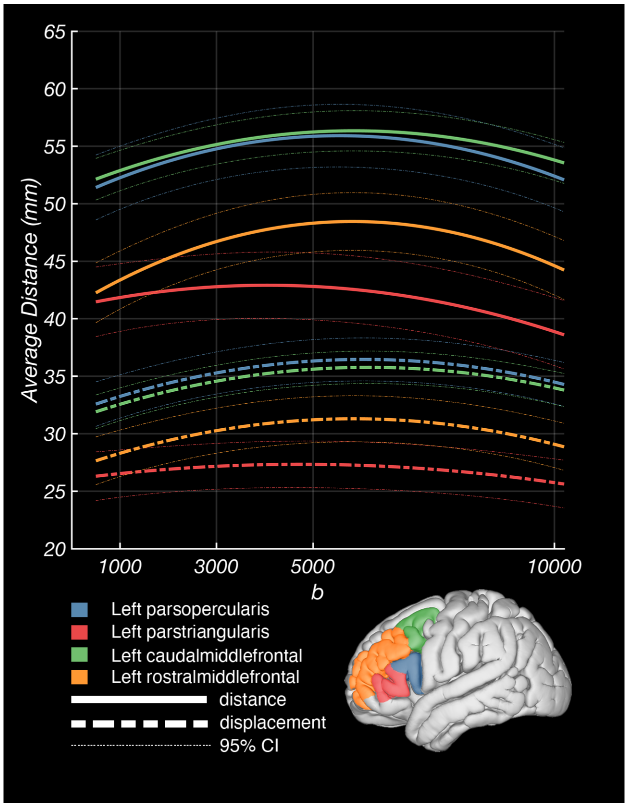


**Figure S2.** Example of distance, displacement, and model fit of four critical language regions in left dorsolateral prefrontal cortex for HCP ultra high **b**-value dataset.
